# RhlR, but not RhlI, allows *P. aeruginosa* bacteria to evade *Drosophila* Tep4-mediated opsonization

**DOI:** 10.1101/165530

**Authors:** Samantha Haller, Adrien Franchet, Abdul Hakkim, Jing Chen, Eliana Drenkard, Shen Yu, Stefanie Schirmeier, Zi Li, Frederick M. Ausubel, Samuel Liégeois, Dominique Ferrandon

**Author notes:** Present address: Department of Pediatrics, Massachusetts General Hospital, Boston, MA 02114 USA. Present address: Institut für Neuro-und Verhaltensbiologie, Badestrasse 9, D-48149 Münster (Stefanie Schirmeier).

## Abstract

When *Drosophila* flies feed on *Pseudomonas aeruginosa* strain PA14, some bacteria cross the intestinal barrier and start proliferating inside the hemocoel. This process is limited by hemocytes through phagocytosis. We have previously shown that the PA14 quorum-sensing regulator RhlR is required for these bacteria to elude the cellular immune response. RhlI synthesizes the auto-inducer signal that activates RhlR. Here, we compare the null mutant phenotypes of *rhlR* and *rhlI* in a variety of infection assays in *Drosophila* and in the nematode *Caenorhabditis elegans*. Surprisingly, in *Drosophila*, unlike *ΔrhlR* mutants, *ΔrhlI* mutants are only modestly attenuated for virulence and are poorly phagocytosed and opsonized in a Thioester-containing Protein4-dependent manner. Likewise, Δ*rhlI* but not Δ*rhlR* mutants colonize the digestive tract of *C. elegans* and kill it as efficiently as wild-type PA14. Thus, RhlR has an RhlI-independent function in eluding detection or counter-acting the action of the immune system. In contrast to the intestinal infection model, *Tep4* mutant flies are more resistant to PA14 in a septic injury model, which also depends on *rhlR*. Thus, the Tep4 putative opsonin can either be protective or detrimental to host defense depending on the infection route.

## INTRODUCTION

*Drosophila melanogaster* is a powerful genetic model organism for the study of innate immunity that has been intensely investigated during the past 25 years (Buchon, Silverman et al., 2014). It thus represents an informative system in which to study host-pathogen interactions using either systemic infection or so-called “natural” infection paradigms, such as oral infection (Bier & Guichard, 2012, Ferrandon, 2013, Igboin, Griffen et al., 2012, Limmer, Quintin et al., 2011b). Genetic analysis has allowed the detailed dissection of the *Drosophila* systemic immune response to microbial infections (Lemaitre & Hoffmann, 2007). In addition to melanization which is mediated by the protease-mediated cleavage of prophenol oxidase into active phenol oxidase (PO), two major NF-kappaB pathways, Toll and Immune deficiency (IMD), regulate the induction of the expression of genes that encode potent antimicrobial peptides, which are active against most bacteria and fungi (Binggeli, Neyen et al., 2014, Dudzic, Kondo et al., 2015, Ferrandon, Imler et al., 2007, Ganesan, Aggarwal et al., 2010, Lemaitre & Hoffmann, 2007). The *Drosophila* systemic immune response is so effective, especially in the case of Gram-negative bacterial infections, that a second arm of host defense, the cellular immune response, has remained comparatively less well studied (El Chamy, Matt et al., 2017, Pean & Dionne, 2014). Indeed, blocking cellular immunity through saturation of the phagocytic apparatus with inert particles does not yield a strong susceptibility phenotype to flies infected by *Escherichia coli*, unless the systemic immune response is at least also partially impaired (Elrod-Erickson, Mishra et al., 2000). Nevertheless, we have found that when two opportunistic pathogens *Serratia marcescens* and *Pseudomonas aeruginosa* are fed to *Drosophila*, the cellular immune response plays a key role in controlling the bacteria that escape from the digestive tract (Limmer, Haller et al., 2011a, Nehme, Liegeois et al., 2007). In both cases, the putative phagocytic receptor Eater plays a crucial role and prevents the development of a rapid bacteremia (Kocks, Cho et al., 2005, Limmer et al., 2011a).

The thioester-containing protein Tep1 opsonizes bacteria in the mosquito species *Anopheles gambiae* (Levashina, Moita et al., 2001). It is unknown whether opsonization also plays a role *in vivo* in *Drosophila*, even though its genome encodes five functional *Tep* loci and a pseudogene (*Tep5*) (Bou Aoun, Hetru et al., 2011). The thioester motif is not present in Tep6 and is therefore thought to be nonfunctional. Indeed, Tep6 is required for the establishment of septate junctions in specific parts of the gut, which explains the lethal phenotype of *Tep6* null mutants (Batz, Forster et al., 2014, Hall, Bone et al., 2014). Tep6 has also been shown to induce autophagy in macrophages via a non cell-autonomous process that involves epithelial cells in which it is expressed (Lin, Rodrigues et al., 2017). A previous study failed to find a role for the remaining *Tep* genes (*Tep1* to *Tep4*) in host defense in several models of bacterial or fungal systemic infections (Bou Aoun et al., 2011), although a study reported that *Tep3* mutant flies are highly susceptible to the nematode parasite *Heterorhabditis bacteriophora* (Arefin, Kucerova et al., 2014). Interestingly, a study led in *Drosophila* cultured S2 cells showed that *Tep2* is required for the phagocytosis of the Gram-negative species *E. coli*, *Tep3* is required for the uptake of the Gram-positive *Staphylococcus aureus*, and unexpectedly that *Tep6* is required to phagocytose the dimorphic yeast *Candida albicans* (Stroschein-Stevenson, Foley et al., 2006).

In contrast to *S. marcescens*, *P. aeruginosa* strain PA14 ultimately manages to establish an exponential infection in the hemocoel four to five days after its ingestion. In a previous study, we showed that a member of the LuxR family of signal receptor-transcriptional regulators in PA14, RhlR, is required to circumvent the cellular immune response (Limmer et al., 2011a). Indeed, *rhlR* mutants are almost avirulent in an intestinal infection model since they remain at very low levels in the hemolymph and kill the infected flies at a much reduced rate. Interestingly, the cellular immune response remains functional until late stages of a PA14 infection, suggesting that hemocytes are not directly targeted by PA14, unlike what happens with *P. aeruginosa* strain CHA, which neutralizes *Drosophila* phagocytosis through the secretion of its ExoS toxin into hemocytes (Avet-Rochex, Bergeret et al., 2005).

RhlR is the major regulator of one of the three known quorum-sensing systems in *P. aeruginosa*. Quorum-sensing systems play a major role in coordinating the expression of virulence genes in several infection models (Coggan & Wolfgang, 2012, Jimenez, Koch et al., 2012, Schuster, Sexton et al., 2013, Williams & Camara, 2009). However, we have failed to uncover a strong role for the other two *P. aeruginosa* quorum sensing systems regulated by LasR and MvfR in the *Drosophila* intestinal infection model (Limmer et al., 2011a). This observation was somewhat unexpected since the LasR-mediated quorum-sensing system appears to function upstream of the RhlR-mediated system. RhlR is activated by binding to an auto-inducer molecule, butanoyl-homoserine lactone (C4-HSL), which is synthesized by the RhlI enzyme. The transcription of the *rhlI* and *rhlR* genes is in turn activated by the Las transcriptional regulator LasR (Coggan & Wolfgang, 2012, Jimenez et al., 2012, Schuster et al., 2013, Williams & Camara, 2009). Activation of RhlR takes place when a threshold concentration of C4-HSL is reached, which correlates with a threshold bacterial concentrations reached during exponential growth in *in vivo* studies.

Here, we report that the virulence phenotype exhibited by ingested PA14 *rhlI* mutants is strikingly distinct from that exhibited by *rhlR* mutants. *rhlR*mutants exhibit severely impaired virulence, whereas *rhlI* mutants at most modestly impaired virulence, suggesting that RhlR can function independently of activation by C4-HSL. We further establish that RhlR, but not RhlI, is required to elude opsonization by a *Tep4*-dependent process. Finally, we establish that in contrast to its protective role during PA14 intestinal infections, Tep4 plays the opposite role in a systemic infection model, possibly by preventing the activation of phenol oxidase.

## RESULTS

### ΔrhlI *is more virulent than* ΔrhlR in the Drosophila *intestinal infection model*

RhlR is activated by C4-HSL synthesized by the RhlI enzyme, as shown in numerous *in vitro* studies (Gambello, Kaye et al., 1993, Latifi, Winson et al., 1995, Pesci, Pearson et al., 1997, Seed, Passador et al., 1995). Thus, we expected that *rhlI* and *rhlR* mutants would display the same phenotype. Unexpectedly, however, although a *ΔrhlI* deletion mutant strain was indeed less virulent than wild-type PA14 when ingested by flies, it was significantly more virulent than *ΔrhlR* (Fig. 1A, B). Moreover, whereas the *ΔrhlR* mutant was cleared from the hemolymph, the *ΔrhlI* mutant proliferated in this compartment, although it appeared to grow less rapidly than wild-type PA14 (Fig. 1C, D). Consistent with this latter result, *ΔrhlI* but not *ΔrhlR* triggered a systemic immune response, as monitored by measuring the expression of the *Diptericin* gene (Fig. S1E-F). Similar results were obtained with independent *ΔrhlR* and *ΔrhlI* in frame deletion mutants constructed by another laboratory (Fig. S1) (Hoyland-Kroghsbo, Paczkowski et al., 2017), thereby confirming the correlation between the *ΔrhlR* and *ΔrhlI* null genotypes and the differing *ΔrhlR* and *ΔrhlI* phenotypes.

**Figure 1:**
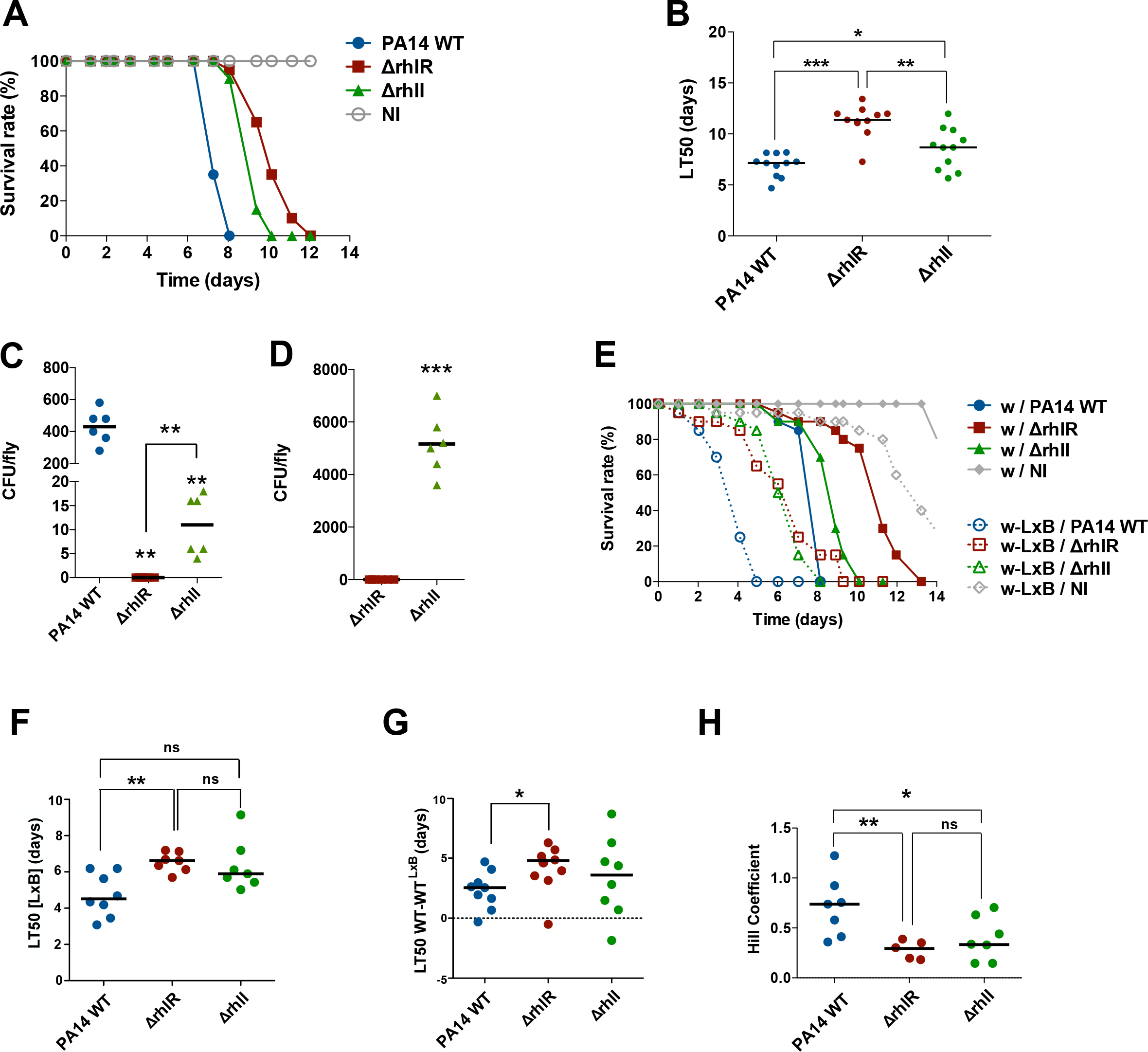
*RhlI* displays a distinct phenotype from that of *rhlR* in the *Drosophila* intestinal infection model. Survival experiment of wild-type flies (*w*) fed on wild-type (WT) PA14 bacteria, or Δ*rhlR*, or Δ*rhlI* mutants. A. Representative survival curves of infected and noninfected (NI) flies. Flies died faster from the infection with PA14 WT than Δ*rhlR* and Δ*rhlI*. Flies infected with Δ*rhlI* exhibited an intermediate survival phenotype. One representative experiment out of seven is shown. Statistical analysis of the data is shown in panel B. B. Pooled LT_50_ data of wild-type flies (*w^A5001^*) following intestinal infections with PA14 WT, Δ*rhlR* or Δ*rhlI*. LT_50_ of flies after infection with PA14 WT was significantly lower than with Δ*rhlR* (***p=0.0003) and ΔrhlI (*p=0.0385). Flies were significantly more susceptible to infection with Δ*rhlI* than with Δ*rhlR* (**p=0.0047). The LT_50_ data from seven survival experiments are displayed (biological duplicates are also shown as there was as much variability between experiments as within experiments), bars indicate medians. Statistics were performed using a non-parametric test (Mann Whitney). C-D. Bacterial titer of the hemolymph collected from flies that had ingested wild-type or mutant PA14 as indicated, three (C) and five (D) days after the ingestion of PA14 WT or mutants as indicated. In this series of experiments, flies infected with PA14 WT had started to succumb by day five and were therefore not analyzed. Statistics were performed using the Mann Whitney test; bars indicate medians. p values for C are respectively from left to right: 0.003, 0.002, and 0.005. P<0.0001 for D. E. Survival curves of wild-type and latex bead-injected flies after intestinal infection with PA14 bacteria. In latex bead-injected flies Δ*rhlI* regains virulence. Note however that the shift in virulence is of the same magnitude as that observed for PA14 WT and contrasts with the large shift observed with Δ*rhlR*. F. Pooled LT_50_ data of latex bead-injected flies (*w*-LXB) survival experiments. *w*-LxB flies died significantly slower after Δ*rhlR* infection than with PA14 WT (**p=0.0065). A slight decrease of virulence, but at the border of significance, was observed between PA14 WT and Δ*rhlI* (p=0.0726). No difference in virulence was detected between Δ*rhlR* and Δ*rhlI* (p=0.3056). Data represent the LT_50_s from five experiments (biological duplicates are also shown as there was as much variability between experiments as within experiments), bars indicate medians. Statistics were performed using a non-parametric test (Mann Whitney). G. Differences of LT_50_s between WT flies pre-injected with PBS (WT) and flies pre-injected with latex-beads (WT^LXB^) after intestinal infection with PA14 WT or Δ*rhlR* mutant or Δ*rhlI* mutants, bars indicate medians. There is only a significant difference between PA14 WT and the Δ*rhlR* mutant (*p=0.0244) but not the Δ*rhlI* mutant. Data represent the LT50s from five experiments (biological duplicates are also shown as there was as much variability between experiments as within experiments), bars indicate medians. Statistics were performed using the Mann Whitney test. H. Hill coefficients of latex bead-injected flies in PA14 infection. Hill coefficients give an indication on the slope of the survival curves. Survival curves of *w*-LXB flies infected with PA14 wild-type (PA14 WT) had a significant higher Hill coefficient than survival curves of flies infected with Δ*rhlR* (**p=0.0092) or Δ*rhlI* (*p=0.0405). No significant difference in Hill coefficient was detected between survival curves of flies infected with Δ*rhlR* or Δ*rhlI* (p=0.6243). The results from three experiments are shown; bars indicate medians. Mann Whitney tests were used for all statistical analyses.

We next tested the survival rates of flies in which the cellular response had been ablated by injecting latex beads (LXB) after feeding on wild-type or *ΔrhlR* or *ΔrhlI* mutant bacteria. Both *ΔrhlR* and *ΔrhlI* killed latex bead-injected flies much faster than PBS-injected control flies, at approximately the same rate, but more slowly than wild-type PA14, with the difference between *ΔrhlI* and wild-type PA14 at the borderline of statistical significance (p=0.07) (Fig. 1E, F).

It is important to determine whether the apparently enhanced virulence of *ΔrhllR* and *ΔrhlI* mutants observed in phagocytosis-impaired flies (compared to flies not injected with latex beads) is simply a reflection of the increased virulence of wild-type PA14 in these immuno deficient flies, or whether the enhanced virulence of the *ΔrhlR* and *ΔrhlI* mutants is indicative of the fact that the RhlR-mediated regulatory systems plays an important role in counteracting the cellular immune response. To this end, it is useful to measure the difference in LT_50_ values of control *vs*. latex bead-injected flies (LT_50_[wt-wt^LXB^]) for each mutant and to compare it to that measured for wild-type PA14. *ΔrhlR* did recover virulence with a LT_50_[wt-wt^LXB^] of 4.7 days, as compared to 2.4 days for wild-type PA14, which corresponds to the level of recovered virulence reported earlier (Fig. 1G) (Limmer et al., 2011a). With a value of 3.5 days, *ΔrhlI* displayed an intermediate LT_50_[wt-wt^LXB^], although the significance of the difference with wild-type PA14 or *ΔrhlR* could not be assessed as the *ΔrhlI* values were too spread out (Fig. 1G). Indeed, *ΔrhlI* mutants consistently tended to display a more variable survival phenotype than the *ΔrhlR* mutant (Fig. 1B).

Finally, both the *ΔrhlR* and *ΔrhlI* mutant strains yielded similarly shaped survival curves when used to infect flies with an impaired cellular defense. These survival curves were less steep than those obtained with wild-type PA14, as measured by their Hill coefficients (Fig. 1H), suggesting that quorum-sensing is involved in determining the shape of the survival curve. This reflects a collective property of flies placed in the same vial, which succumb less synchronously during infections when the *rhl* quorum sensing system is missing.

### rhlI *and wild-type PA14, but not* rhlR, *strains colonize the* C. elegans *digestive tract*

The nematode *C. elegans* is a well-established model host to study *P. aeruginosa* pathogenesis (Irazoqui, Urbach et al., 2010, Pukkila-Worley & Ausubel, 2012, Tan, Mahajan-Miklos et al., 1999a). The *C. elegans* intestinal infection model shares some key features with the *Drosophila* model, at least during the initial stages of the infection. For instance, in both models, ingested bacteria are exposed in the gut lumen to antimicrobial peptides and to reactive oxygen species generated by the Dual oxidase enzyme {Ha, 2005 #1857}{Chavez, 2009 #3616}. However, in contrast to ingested *P. aeruginosa* in *Drosophila*, PA14 are not known to escape from the gut compartment during the nematode infection. We therefore tested whether the differences that we observed in the virulence of *ΔrhlR* and *ΔrhlI* mutants in the *Drosophila* intestinal infection assay were reflected in a well-established *C. elegans – P. aeruginosa* nematode “slow killing” survival assay (McEwan, Feinbaum et al., 2016, Tan et al., 1999a). Indeed, two independently constructed in frame *rhlR* deletion mutants in the PA14 background were dramatically less virulent than two independently constructed *rhlI* deletion mutants in their ability to kill *C. elegans* (Fig. 2A)

**Figure 2:**
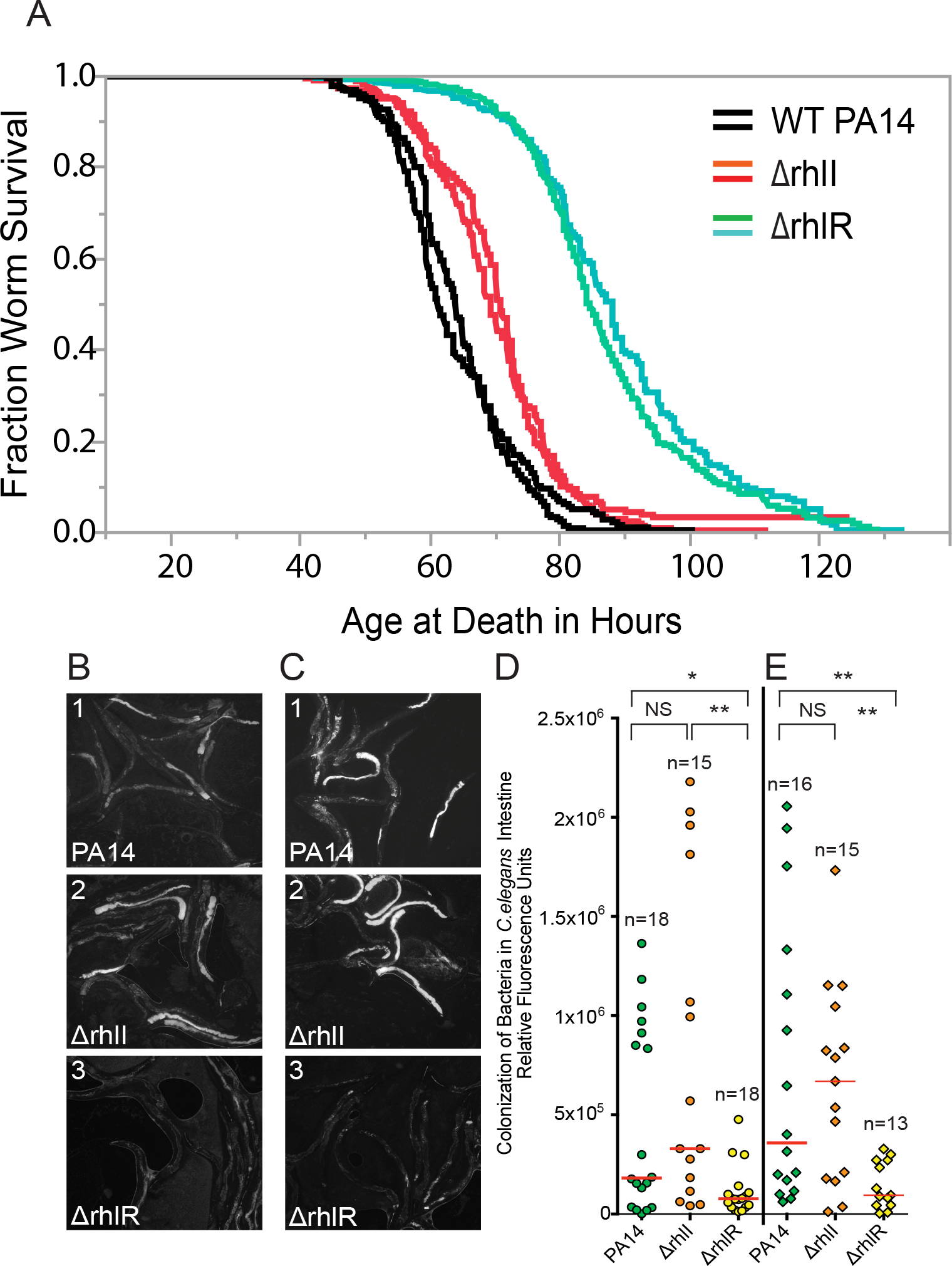
*rhlR* mutants are impaired in their ability to kill *C. elegans* and do not colonize the *C. elegans* intestine. A, an automated *C. elegans* life-span machine was used to monitor the survival of worms in a *P. aeruginosa*-mediated killing “slow killing” assay (McEwan et al., 2016). Approximately 200 wild-type *C. elegans* nematodes were fed WT PA14, or *ΔrhlI*, or *ΔrhlR PA14* mutants (constructed in the Ausubel laboratory (red (*ΔrhlI)* and blue *(ΔrhlR)*) or Bassler (orange (*ΔrhlI)* and green *(ΔrhlR)*) laboratories. P<0.001 (log-rank test) for PA14 vs *ΔrhlI*, PA14 vs *ΔrhlR, or ΔrhlI vs ΔrhlR*. The experiment was repeated twice with similar results. B-E, *C. elegans* wild-type N2 animals were fed WT PA14, *ΔrhlI*, or *ΔrhlR PA14* mutants (constructed in the Ausubel laboratory (B and D) or Bassler laboratories (C and E) expressing GFP. At 48 h post infection, 13-18 worms infected with WT PA14 or the *ΔrhlI* or *ΔrhlR* mutants were imaged in the green fluorescent channel. B and C. Representative images are shown. D and E. Images were quantified using ImageJ. There was a significant difference in the levels of live bacteria between WT PA14 and *ΔrhlR* (p = 0.034 (D) and 0.0097 (E)), and there were significant differences between the *ΔrhlR* and *ΔrhlI* mutants (p = 0.0012 (D) and 0.0042 (E)), using a non-parametric Mann-Whitney test. The differences between *ΔrhlI* and WT PA14 were not significant (p = 0.16 (D) and p = 0.95 (E)). The experiments were repeated at least two times with similar results.

The primary *C. elegans* immune response occurs in intestinal epithelial cells and because the worms are transparent, host-pathogen interactions can be easily visualized in the intestinal lumen. Thus, in addition to the *C. elegans* survival assay (Fig. 2A), we also used a quantitative assay (Figs. 2B-E; see Materials and Methods) that monitors the accumulation of live bacterial cells in the intestine of the nematodes using PA14, *ΔrhlR*, and *ΔrhlI* expressing GFP to monitor the level of intestinal colonization. Live wild-type *P. aeruginosa* PA14 cells start accumulating in the intestine 24-48 hours post infection. Consistent with the *Drosophila* infection assays described in Fig. 1, two independent *ΔrhlR* mutants in the PA14 background colonized the *C. elegans* intestine at significantly lower levels than two independent *ΔrhlI* mutants. Indeed, in this colonization assay, the *ΔrhlI* mutants were indistinguishable from wild-type PA14, similar to the results in the nematode killing assay. An alternative explanation for these results is that *C. elegans* preferentially feeds on the *ΔrhlI* mutant compared to the *ΔrhlR* mutant and simply overwhelms the immune response with a larger number of ingested cells. However, this possibility was ruled out by monitoring the pumping (feeding) rate of *C. elegans* feeding on wild-type, *ΔrhlR* and *ΔrhlI* mutants. *C. elegans* pumped at the same rate on all three strains (Fig. S2).

### *Phagocytosis protects* Drosophila *against invasion of its hemocoel by wild-type PA14 during the early phase of the infection*

In the *Drosophila* intestinal infection model, flies constantly feed on PA14 present on a filter. A defining feature of this model is that even though bacteria are able to rapidly cross the intestinal barrier to penetrate the hemocoel, as had been previously described for *Serratia marcescens* (Nehme et al., 2007), the PA14 titer in the hemolymph remains low for the first few days of the infection. After this initial incubation period, there is an exponential proliferation of PA14 in the hemocoel, which coincides with the activation of the systemic immune response. Using bacteria expressing different colored fluorescent proteins, we previously showed that *S. marcescens* continuously crosses the intestinal barrier during the infection (Nehme et al., 2007). Fig. 3A-B shows that when flies that have been feeding on dsRed-labeled PA14 bacteria for four days were switched to a filter laced with GFP-labeled PA14, the green bacteria progressively replaced the red bacteria both in the gut and hemocoel compartments. We conclude that *P. aeruginosa*, like *S. marcescens*, continuously crosses the intestinl barrier during the infection.

**Figure 3:**
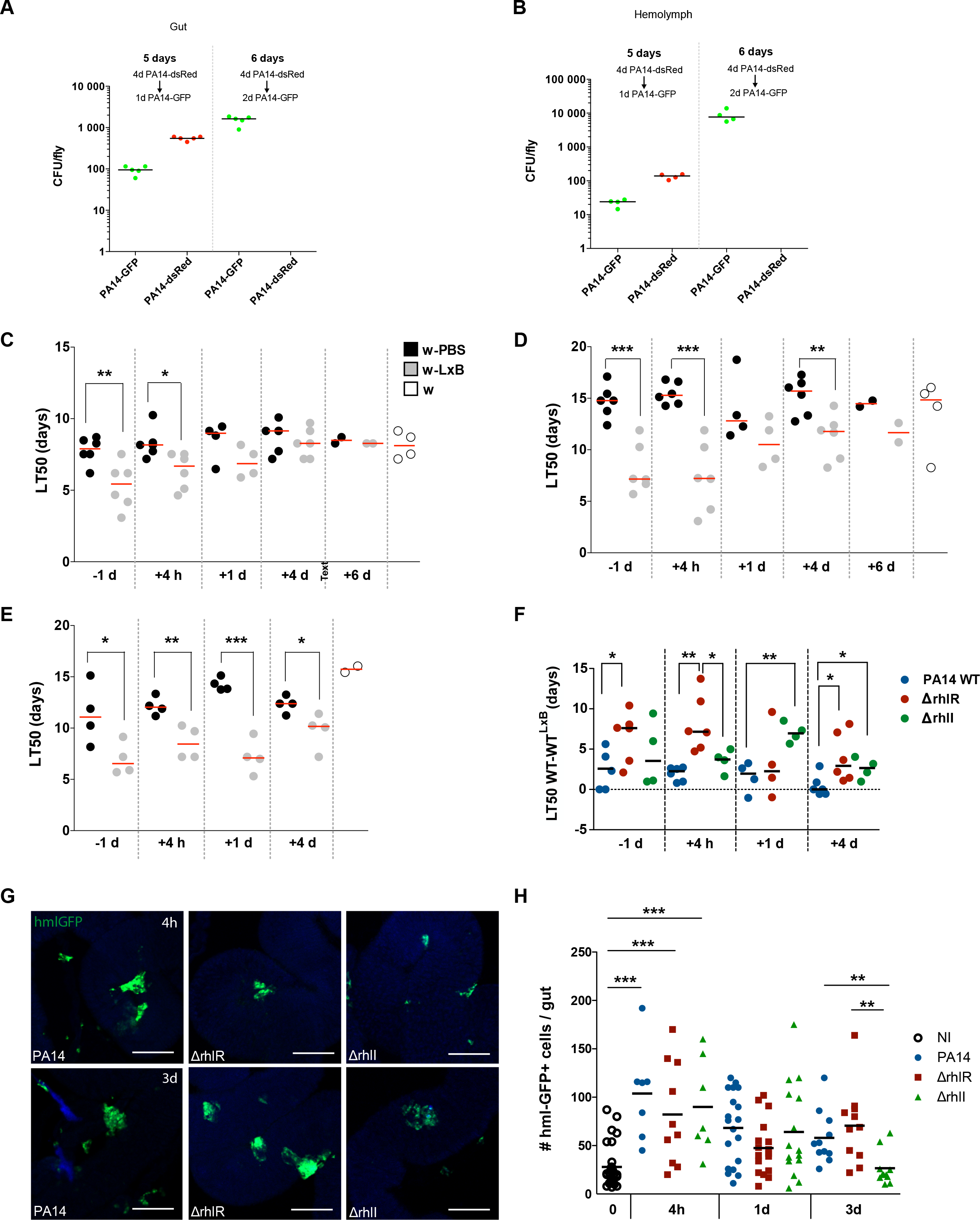
Phagocytosis is required during the early stage of the infection in *Drosophila*. A and B, Wild-type *Drosophila* were orally infected with wild-type PA14 expressing dsRed (PA14-dsRed). After 4 days infected flies were transferred to tubes containing wild-type PA14 expressing GFP (PA14-GFP). At 5 days of the infection (one day after transferring flies to PA14-GFP), most PA14 bacteria in the *Drosophila* gut expressed dsRed and only 10% expressed GFP (A). However, at 6 days (2 days after the transfer of flies to PA14-GFP), only GFP positive bacteria were detected in the gut. Results were similar for bacteria retrieved from the hemolymph compartment (B). C, D and E, LT_50_s from survival experiments of WT flies (w) after intestinal infection with PA14 WT (C), Δ*rhlR* (D) or Δ*rhlI* (E) mutants and injection of either latex beads (LXB, grey dots) or PBS (black dots) at different time points of the infection. Latex beads or PBS were injected either one day before the infection started (-1d) or four hours (+4h), one day (+1d), four days (+4d) or six days (+6d) after the infection started. White dots correspond to the survival of infected, uninjected flies. (C) LT50s of *w^A5001^*-LxB are significantly lower than *w^A5001^* only at -1d (**p=0.0086) and +4h (*p=0.0154). (D) LT_50_s of *w*-LXB flies are significantly lower than *w* at most times during the infection (-1d: ***p=0.0002, +4h: ***p=0.0002 and +4d: **p=0.0069). (E) A similar phenotype is observed with flies infected with Δ*rhlI* (-1d: *p=0.0395, +4h: **p=0.0085, +1d: ***p=0.0002 and +4d: *p=0.0400). Note however that for injections of latex beads at day4 the difference is reduced, as compared to earlier time points of injection of latex-beads. The cumulative LT_50_ data from at least three experiments (only two experiments for Δ*rhlI*) are shown, except for +6 d; bars indicate medians. Statistical analyses were done with an unpaired t-test. F. Differences of LT_50_s between WT flies injected with PBS (WT) and flies injected with latex-beads (WT^LXB^) after intestinal infection with PA14 WT or Δ*rhlR* mutant or Δ*rhlI* mutants in at least two experiments, bars indicate medians. G. Guts of transgenic flies with GFP-labeled hemocytes were dissected in a manner that preserves the association of hemocytes with the digestive tract and were examined by fluorescence confocal microscopy. Green: GFP; blue: DAPI staining of nuclei. Scale bars: 100µM. H. Analysis of hemocytes recruited to the fly intestine upon the infection with either wildtype PA14, or Δ*rhlR* or Δ*rhlI* mutant bacteria. Both PA14 WT and the Δ*rhlR* or Δ*rhlI* mutant bacteria induce a recruitment of hemocytes to the gut (4h after the beginning of the infection, for each bacteria p<0.0001). At one day, there are slightly more hemocytes recruited when infected with wild-type PA14 compared to the mutant. While at 3 days after the beginning of the infection there are more hemocytes recruited when infected with the Δ*rhlR* mutant bacteria (p=0.0025 between Δ*rhlR* and Δ*rhlI*). Data represent 3 pooled experiments. Bars indicate medians.

Next, we asked at what time periods during an infection is phagocytosis important in preventing PA14 growth in the hemolymph. To this end, we saturated the phagocytic apparatus of hemocytes by injecting latex beads into flies at different time points during infection. As expected, blocking phagocytosis one day prior to the infection led to an earlier demise of the PA14-infected flies compared to PBS-injected control flies. Similar results were found when latex beads were injected four hours or one day after infection, although in the latter case the difference was not significant (its value was nevertheless similar to that obtained by injection one day prior to infection at -1 day; Fig. 3C). In contrast to injecting the latex beads one day after infection, the injection of latex beads four or six days after the beginning of the ingestion of wild-type PA14 did not modify the survival rate of flies. That is, the LT_50_ values were similar to those of control (PBS-injected) flies, consistent with the conclusion that starting about four days after infection the cellular immune response no longer plays a major role in limiting a wild-type PA14 infection.

In contrast to wild-type PA14, *ΔrhlR* bacteria were kept in check by phagocytosis at least up to day four and to some extent up to six days after infection (Fig. 3D, F). Phagocytosis was efficient against *ΔrhlI* bacteria for approximately four days (Fig. 3E, F). These data suggest that *ΔrhlR* bacteria are constantly kept in check by the cellular immune response when penetrating the hemocoel, whereas wild-type PA14 ultimately escape this immune surveillance. *ΔrhlI* bacteria display an intermediate phenotype, with a somewhat decreased virulence in flies in which phagocytosis was blocked at day 4 (Fig. 3F).

A recent study has reported that hemocytes are recruited to the gut after the ingestion of bacteria (Ayyaz, Li et al., 2015). We confirmed this finding in the case of *P. aeruginosa* oral infection, with a significant recruitment observed at four hours after the beginning of the infection with either wild-type PA14 or *Δrhl* mutants (Fig. 3G-H, Fig. S3). While hemocytes remained associated with the midgut for at least three days after the beginning of the ingestion of wild-type PA14 or *ΔrhlR* bacteria, they were not at three days in the case of *ΔrhlI* bacteria (Fig. 3H). While some ingested bacteria could be detected in the hemocytes recruited to the gut, this phenomenon was not reproducible enough to allow reliable quantification.

### Drosophila Tep4 *is required for host defense against ingested PA14*

Our previous work had shown that flies mutant for the putative phagocytic receptor gene *Eater* are more susceptible to PA14 ingestion and display a phenotype similar to that obtained by latex bead-mediated ablation of the phagocytic capacity of hemocytes (Limmer et al., 2011a). Thioester-containing proteins have been reported to be required for phagocytosis in mosquitoes and also in cultured *Drosophila* S2 cells (Levashina et al., 2001, Stroschein-Stevenson et al., 2006). We therefore tested mutations affecting the *Tep2*, *Tep3*, and/or *Tep4* genes (Bou Aoun et al., 2011). In the case of *Tep1*, since no mutants were available, we tested an RNAi transgene expressed either in hemocytes or in the fat body. However, we did not observe any change in the virulence of ingested PA14 (Fig. S4A, B). *Tep4* and triple *Tep2-Tep3-Tep4* mutants displayed increased susceptibility to PA14 ingestion, in contrast to *Tep3* and double *Tep2-Tep3* mutants that displayed respectively a somewhat decreased or wild-type susceptibility (Fig. 4A, D-E). Of note, uninfected *Tep3* mutants fed on a sucrose solution displayed an enhanced fitness when compared to wild-type or other *Tep* mutant lines (Fig. S4C). We conclude from these data that Tep4, but not other thioester-containing proteins, is required for host defense against ingested PA14.

**Figure 4:**
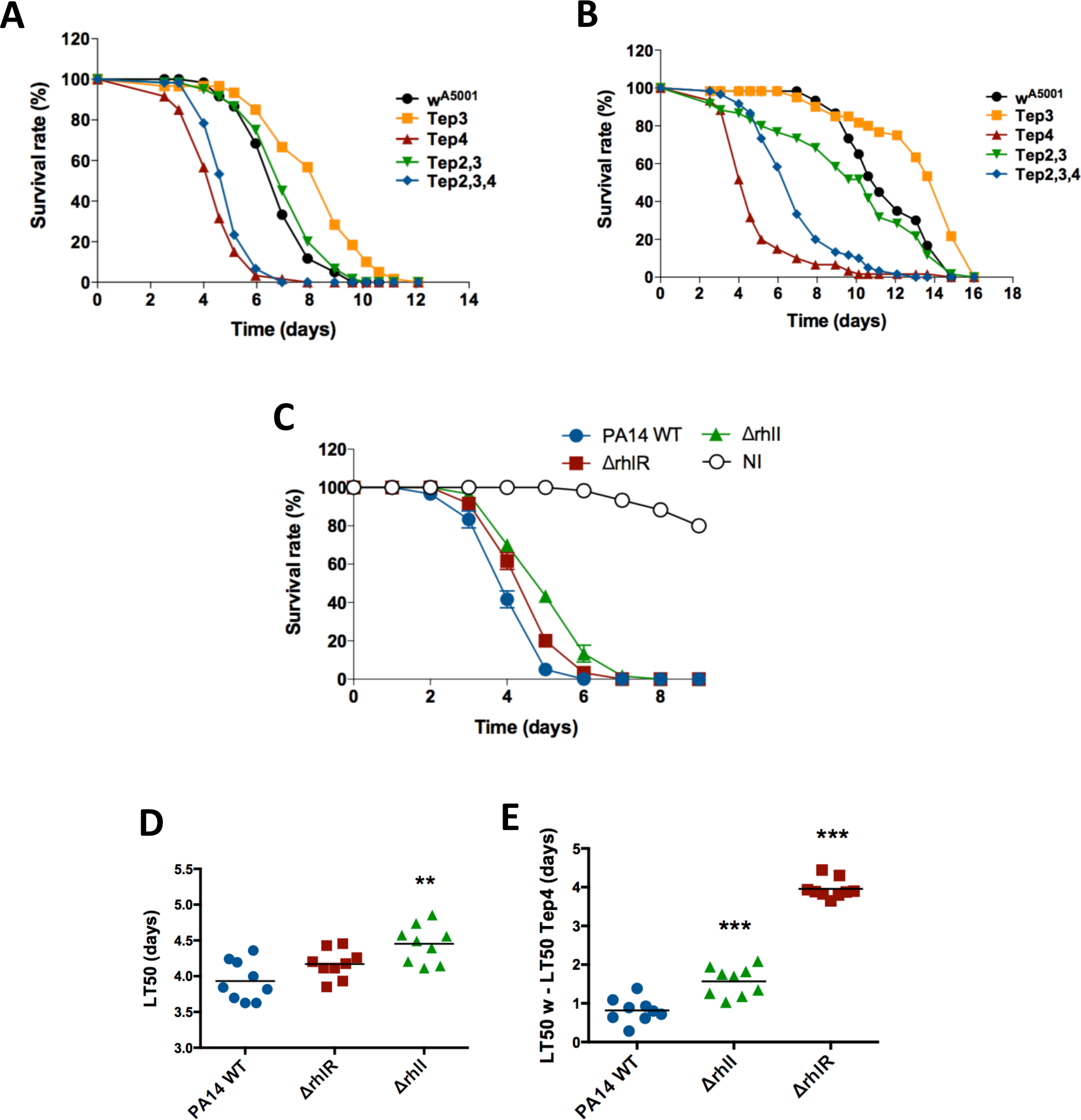
*RhlR* circumvents *Tep4*-mediated host defense. A and B. *Drosophila* wild-type flies (*w^A5001^*), single mutants *Tep3* and *Tep4*, double mutant *Tep2,3* and triple mutant *Tep2,3,4* were orally infected with PA14 WT (A) or the Δ*rhlR* mutant (B) in parallel survival experiments. A. *Tep4* and *Tep2,3,4* mutant flies are significantly more susceptible to PA14 infection compared to *w^A5001^*. No difference in survival was detected between the *Tep2,3* mutant and *w^A5001^*. Surprisingly *Tep3* mutants seemed to be more resistant to infection. B, A strong enhancement of Δ*rhlR* virulence is observed with *Tep4* and *Tep2,3,4* mutants compared to *w^A5001^* flies. *Tep2,3* and *w^A5001^* exhibited nearly the same rate of death when challenged with Δ*rhlR*. The *Tep3* mutant seemed again to be more resistant to the infection. In A and B, one representative experiment out of three (each with biological triplicates, except for uninfected flies) is shown. C. The survival of *Tep4* flies infected with PA14 WT or Δ*rhlR* or Δ*rhlI* bacteria was examined. One representative experiment out of three (each with biological triplicates) is shown. D. Quantification of the experiments shown in C. The triplicates were assessed as independent experiments as there was as much variability between experiments as within experiments. p values comparing the LT_50_s of mutants versus PA14 WT are from left to right: 0.07, **: 0.003, (Mann-Whitney test). Bars indicate medians. E. Difference of LT_50_s between *w* and *Tep4* flies after intestinal infection with either PA14 WT, Δ*rhlR*, or Δ*rhlI* bacteria. p values comparing the LT_50_s of mutants versus PA14 WT : *** : p<0.0001 (Mann-Whitney test). Bars indicate medians.

Next, we found that *ΔrhlR* became as virulent as wild-type PA14 when ingested by *Tep4* or *Tep2-Tep3-Tep4* mutants, which was not observed with the *Tep2* and *Tep2-Tep3* mutant strains (Fig. 4B, D-E). Interestingly, the injection of latex beads in *Tep4* flies only modestly increased the virulence of *rhlR* bacteria when compared to PBS-injected *Tep4* flies (Fig. S4 D, E), suggesting that phagocytosis of PA14 is severely affected in the *Tep4* mutant. *ΔrhlI* bacteria behaved like *ΔrhlR* bacteria when ingested by *Tep4* (Fig. 4C, E), similarly to flies injected with latex beads (Fig. 1E), although *ΔrhlR* recovered virulence to a much larger extent (3.1 days) than *ΔrhlI* (0.9 days) when ingested by *Tep4* flies. Hence, the behavior of *ΔrhlR* mutant PA14 is similar in *eater* and *Tep4* mutant flies, thereby raising the possibility that both fly genes are involved in the same process, in keeping with an unchanged phenotype of *Tep4* when phagocytosis was blocked by the injection of latex beads (Fig. S4DC).

### *Phagocytosis of PA14 bacteria is impaired in* Tep4 *mutant hemocytes*

To quantitatively monitor the uptake of PA14, we used assays that relied on larval hemocytes. First, we injected heat-killed, pHrodo®-labeled bacteria in wild-type or *Tep4* third-instar larvae. The dye becomes fluorescent when placed in an acidic environment such as that encountered in phagosomes. After 45 minutes, the larvae were bled and a phagocytic index was established. Wild-type hemocytes ingested significantly more PA14 or *ΔrhlR* bacteria than *Tep4* hemocytes (Fig. 5A). There were however no significant differences between heat-killed wild-type PA14 and *rhlR* bacteria uptake by wild-type hemocytes on the one hand, or *Tep4* hemocytes on the other (Fig. 5A). This was not necessarily unexpected as heat-killing likely inactivates *rhlR*-dependent virulence factors and might also alter the surface of bacteria. We therefore modified the assay with live bacteria and used an antibody we had raised against PA14 to differentially immuno-stain the bacteria, prior to and after permeabilization of the fixed cells. As before, *Tep4* hemocytes exhibited a decreased uptake of bacteria compared to wild-type hemocytes. However, in the case of live bacteria, *ΔrhlR* bacteria were significantly better phagocytosed than wild-type PA14 bacteria when injected into wild-type or *Tep4* larvae. *ΔrhlI* exhibited an intermediate phenotype in this assay and was not significantly different from either wild-type or *ΔrhlR* bacteria (Fig. 5B). We conclude that this assay is not sensitive enough to discriminate between these two bacterial mutant strains. We obtained similar results using the *ΔrhlR* and *ΔrhlI* strains generated by another laboratory (Fig. S1G).

**Figure 5:**
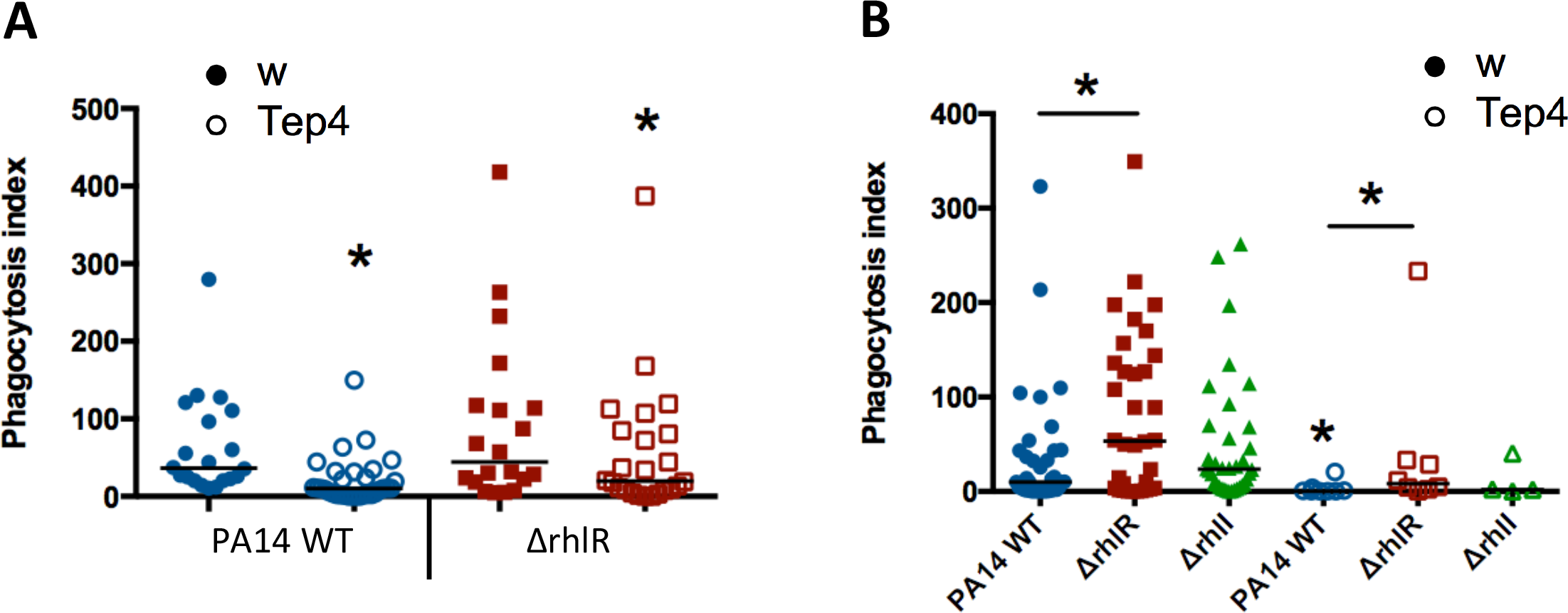
Tep4 is required for phagocytosis of *rhlR* mutant bacteria. A. Heat-killed pHrodo®-labeled bacteria of the indicated genotype were injected into either wild-type or *Tep4* third instar larvae and incubated for 45 minutes. The hemocytes were then retrieved. Bacteria present in phagosomes were fluorescent and used to measure the phagocytic index. Bacteria injected into *Tep4* were significantly less phagocytosed than those injected into wild-type larvae: p<0.0001 for PA14 WT and p=0.04 for Δ*rhlR* (Mann-Whitney test). Bars indicate medians. B. The experiment is similar to that shown in A, except that live bacteria were used and a differential antibody staining protein was used to reveal phagocytosed bacteria. PA14 WT, Δ*rhlR*, Δ*rhlI* bacteria were injected in wild-type or *Tep4* larvae. Δ*rhlR* bacteria were more readily phagocytosed than PA14 WT both by wild-type (p=0.01) or *Tep4* (p=0.04) hemocytes (Mann-Whitney test). Bars indicate medians.

### *Tep4 opsonizes* rhlR *better than* rhlI *or wild-type bacteria*

We next designed an experiment to assess whether Tep4 functions as an opsonin, *i.e*., that it is deposited on the surface of bacteria to facilitate its detection and subsequent ingestion by hemocytes. To this end, we collected the hemolymph from either wild-type or *Tep4* larvae and incubated it with live bacteria. These bacteria were then retrieved and injected into either naive wild-type or *Tep4* mutant larvae prior to bleeding these injected larvae to be able to count the ingested bacteria as described above (Fig. 6A). Wild-type PA14 were poorly phagocytosed in this assay (medians of phagocytic index lower than 10, Fig. 6C), when the opsonized bacteria were first incubated in the hemolymph of Tep4-containing wild-type larvae and then secondarily injected into wild-type or *Tep4* larvae (although the former yielded a significantly increased phagocytic index, as compared to bacteria incubated first in *Tep4* hemolymph; Fig. 6C). In contrast, bacteria that had been first incubated in *Tep4* mutant hemolymph were hardly ingested when re-injected into *Tep4* larvae (Fig. 6E). Re-injection of these bacteria into wild-type larvae recipients modestly increased the phagocytic index (Fig. 6E), which was nevertheless lower than that of bacteria that had been pre-incubated with wild-type hemolymph (Fig. 6C). When these experiments were performed using *ΔrhlR* bacteria that were injected in *Tep4* larvae, it made a major difference whether these mutant bacteria had first been pre-exposed to wild-type or *Tep4* hemolymph. Tep4-dependent opsonization led to a massive uptake of bacteria (median phagocytic value of 162), whereas nonopsonized bacteria (pre-incubated with *Tep4* mutant hemolymph) were hardly ingested (median phagocytic value of 12; Fig. 6D-E). As expected, nonopsonized bacteria that were then injected in wild-type recipients were much better phagocytosed (median phagocytic value of 214), presumably because Tep4 was circulating in the wild-type hemolymph (Fig. 6B). They were nevertheless ingested less efficiently than opsonized bacteria (median phagocytic value of 376; Fig. 6B). Finally, *ΔrhlI* bacteria were also opsonized by Tep4, but significantly less than *ΔrhlR* bacteria (Fig. 6B, D). Again, they displayed a phenotype that was intermediate to that of wild-type PA14 on the one hand, and *ΔrhlR* on the other. We conclude that PA14 and, to a lesser extent, *ΔrhlI* bacteria are less efficiently opsonized and subsequently phagocytosed than *ΔrhlR* bacteria, which are therefore unable to elude the cellular immune response.

**Figure 6:**
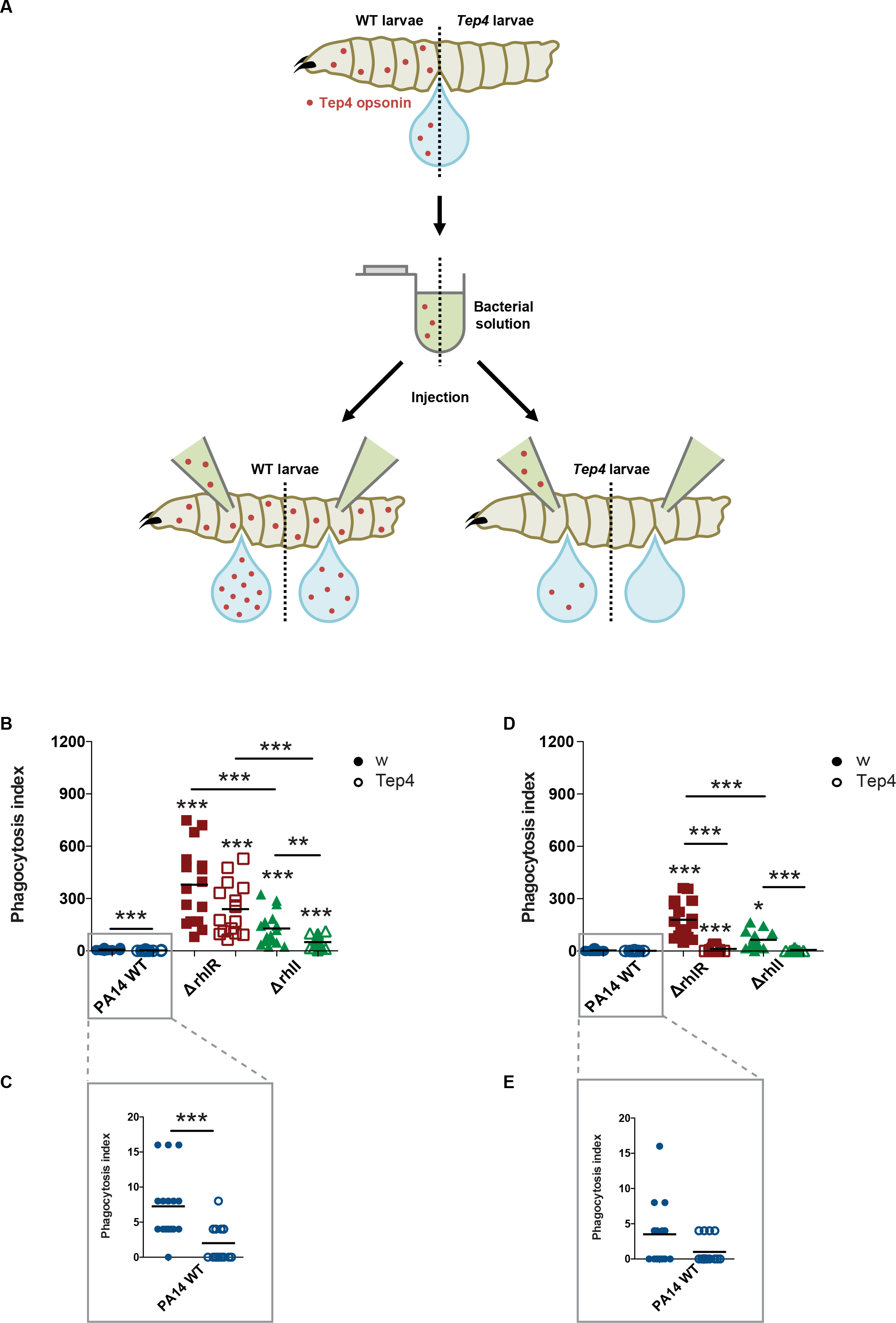
Tep4 is an opsonin that preferentially detects *rhlR* over *rhlI* mutant bacteria. A. Scheme of the experimental procedure. Live bacteria were incubated with either wild-type or *Tep4* hemolymph and were thereafter injected into naive larvae, which were either wildtype or *Tep4*. The phagocytosis index was then measured as in Fig. 6B. B-E. Bacteria pre-incubated with wild-type or *Tep4* hemolymph are represented in pairs, respectively with filled (left) and open (right) circles. Bars indicate medians. Data were analyzed using the Mann-Whitney test. ***: p<0.0001 B. PA14 WT, Δ*rhlR*, and Δ*rhlI* bacteria pre-incubated with either wild-type (wt) or *Tep4* hemolymph were injected into wild-type larvae. Stars above symbols report the statistical significance of the data when compared to PA14 WT. **: p=0.007 D. PA14 WT, Δ*rhlR*, and Δ*rhlI* bacteria pre-incubated with either wild-type or *Tep4* hemolymph were injected into *Tep4* larvae. **: p=0.01 C, E. These panels display respectively a magnification of B and D to show the low phagocytic index associated with PA14 WT uptake.

### *Tep4 plays an adverse role in a PA14 systemic infection model in* Drosophila

A recent study has reported that *Tep4* mutants are more resistant to a systemic infection with the entomopathogenic bacterium *Photorhabdus luminescens* in a septic injury model (Shokal & Eleftherianos, 2017). By injecting several doses of PA14, from 10 to 1000 CFUs, directly into the thorax of flies, we also consistently found that *Tep4* mutants survived better than wild-type flies in this systemic infection model (Fig. 7A). This difference in survival between *Tep4* and wild-type flies was largely attenuated when *ΔrhlR* bacteria were injected, thereby establishing again a relationship of altered virulence of these bacteria in a *Drosophila Tep4* mutant background (Fig. 7B). Using the steady-state expression of the antibacterial peptide gene *Diptericin* as a read-out of the activation of the IMD pathway that regulates the systemic immune response against Gram-negative bacteria, we found no significant difference of expression between wild-type and *Tep4* (Fig. 7C). As a higher level of induction of the IMD pathway is unlikely to account for the increased resistance of *Tep4* mutants against PA14 infection, we tested whether the phenol oxidase cascade was more efficiently activated in this mutant background, as previously reported (Shokal & Eleftherianos, 2017). Indeed, we found that pro-phenol oxidase was cleaved to some extent in *Tep4* but not in wild-type flies. These data further suggest that wild-type PA14 bacteria elude detection by the factors that trigger the prophenol oxidase cascade, and that Tep4 plays a role in this process of evasion from the melanization response.

**Figure 7:**
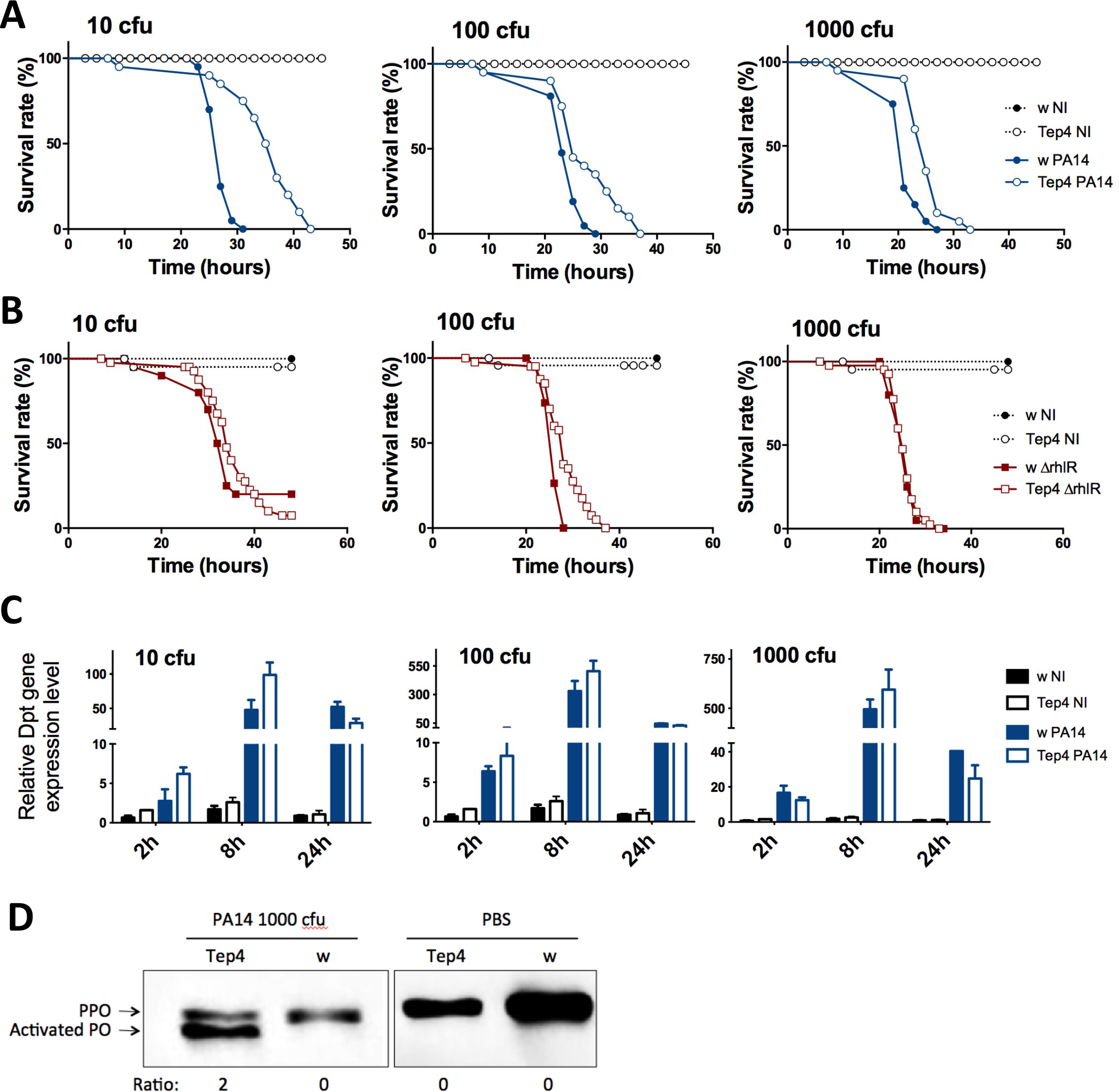
*Tep4* mutants are more resistant to a systemic infection with PA14. A and B. The survival of wild-type and *Tep4* flies was examined, after injection of PBS as a non-infected control (NI), or 10, 100, 1000 cfus of PA14 WT (A) or PA14 Δ*rhlR* (B) bacteria.C. *Diptericin* expression was measured by RT-qPCR in wild-type and *Tep4* flies, 2, 8 or 24h after injection of PBS (NI), or 10, 100, 1000 cfus of PA14 WT bacteria. Each experiment was performed independently three times and a representative experiment is shown (A-C). The *Diptericin* expression shown is relative to *RpL32* expression and normalized to the noninfected wild-type control. D. The cleavage of the prophenol oxidase have been analyzed 2 hours after injection of 1000 cfus of PA14 WT bacteria or PBS, by Western-blotting using a pan-phenol oxidase antibody. The intensity of the prophenol oxidase (PPO) and of the cleaved active phenol oxidase (activated PO) bands was measured and the ratio of the measurements is shown below the blot. This experiment was performed twice with a similar result.

## DISCUSSION

In this study, we analyzed the interactions of *P. aeruginosa* with *Drosophila* from the dual perspective of both pathogen and host. Our data lead us to propose a model in which RhlR plays a pivotal role in virulence by diminishing the ability of the cellular immune arm of the host defense response to detect *P. aeruginosa* once the bacteria have reached the internal body cavity of the insect after crossing the intestinal barrier. Surprisingly, RhlR function in eluding opsonization by Tep4 is at least partially independent of the C4-HSL producing enzyme RhlI. These results as well as those of another study (Mukherjee, Moustafa et al., 2017) show that RhlR can function independently of C4-HSL, but they do not formally establish that RhlR is functioning independently from a quorum-sensing system in this function. Furthermore, we establish a dual function for the putative opsonin Tep4, which plays opposite roles in host defense depending on the infection route.

### *A* rhlI-*independent function of* rhlR

*P. aeruginosa* is a pathogen that uses complex signaling mechanisms to adapt to its environment, and its three quorum-sensing regulators (LasR, RhlR and MvfR) appear to be involved in the regulation of a variety of virulence-related functions (Coggan & Wolfgang, 2012, Jimenez et al., 2012, Schuster et al., 2013). *In vitro* studies, sometimes complemented by *in vivo* experiments, have revealed that these quorum-sensing systems are intricately intertwined. It was thus unexpected that only the *P. aeruginosa* RhlR regulator appears to play a critical role for virulence in a *Drosophila* intestinal infection model and not the two other quorum sensor regulators (Limmer et al., 2011a). Here, we report that *ΔrhlR* null mutants consistently display virulence levels that are much weaker than those observed with *ΔrhlI* mutants (Fig. 1B), do not proliferate in the hemolymph in contrast to *ΔrhlI* mutants, and are phagocytosed and opsonized in a Tep4-dependent process more efficiently than *ΔrhlI* and wild-type bacteria. These observations suggest that RhlR functions at least partially independently from RhlI and presumably independently from C4-HSL activation. In contrast, *ΔrhlI* mutants exhibit an impaired virulence in both wild-type and *Tep4* immuno-deficient flies, similarly to *ΔrhlR* mutants. Both *ΔrhlI* and *ΔrhlR* mutants display survival curves with shallow slopes. Thus, the partially overlapping phenotypes of *ΔrhlI* and *ΔrhlR* in flies with impaired cellular immunity suggests that RhlR partially functions together with RhlI as a classical quorum-sensing regulator in a process that remains to be delineated. Interestingly, it appears that RhlR controls gene expression for biofilm formation both in a C4-HSL-dependent and C4-HSL independent manner (Mukherjee et al., 2017). However, this putative conventional function of RhlR plays a minor role in virulence of PA14 in *Drosophila*, as shown by the weak virulence-related phenotypes of *ΔrhlI* mutants documented in our study.

One explanation for the RhlI-independent activity of RhlR is that it gets activated in a C4-HSL-independent manner by an as yet unidentified quorum-sensing compound. Of note, RhlR does not appear to be activated by 3-oxo-C12-HSL (Mukherjee et al., 2017), the LasR ligand. In any case, *lasR* and *lasI* mutant bacteria display only a modestly decreased virulence phenotype in the *Drosophila* infection model (Fig. S5). The diketopiperazines (DKPs) represent a candidate family of RhlR-activating compounds (Holden, Ram Chhabra et al., 1999); however, at least one study failed to detect any interaction of these compounds with LuxR proteins (Campbell, Lin et al., 2009). The resolution of this issue will require testing mutants that affect the synthesis of DKPs.

Another hypothesis to consider is that RhlR may function independently of auto-inducer molecules. RhlR forms dimers in the presence or absence of C4-HSL (Ledgham, Ventre et al., 2003). Further studies reported that RhlR functions as a repressor when unbound to C4-HSL (Anderson, Zimprich et al., 1999, Medina, Juarez et al., 2003). Interestingly, RhlR dimers seem to bind its target DNA sequence with an altered conformation (Medina et al., 2003). Finally, transcriptomics studies on *lasR-rhlR* double mutants also revealed several target genes that appear to be repressed by either LasR or RhlR (Schuster, Lostroh et al., 2003, Wagner, Bushnell et al., 2003). Thus, a repressor function for RhlR unbound to C4-HSL cannot be excluded at this stage. A limitation of all of the studies related to the C4-HSL-related activity of RhlR is that they were performed *in vitro* and not *in vivo*.

Finally, our studies on the inactivation of the cellular immune response at different time points of the infection further support a quorum sensing-independent role of RhlR. Our study revealed that phagocytosis is required only when very few bacteria are present in the hemolymph, that is, during the first days of the infection. A caveat here is that we cannot exclude the possibility that C4-HSL or other cryptic autoinducers might be produced by the bacteria present in large numbers in the gut compartment. However, if autoinducers, including C4-HSL, were produced in the intestinal lumen and were able to cross the digestive barrier, it is difficult to understand why they would not immediately activate RhlR resulting in full-blown bacteremia without the observed lag before the exponential proliferation phase in the hemocoel. This hypothesis also does not account for why the *rhlI* reduced virulence phenotype is much weaker than that of *rhlR*, unless this reflects the differing opsonization properties of these mutants.

### *The function of RhlR in bypassing host defenses in* C. elegans?

Except for fungal invasion of the epidermis by nematophagous fungi, how *C. elegans* senses infections remains poorly understood (Zugasti, Bose et al., 2014). The finding that *ΔrhlR* mutants are much less virulent than *ΔrhlI* mutants in a *C. elegans* infection model, and that *ΔrhlR* mutants fail to colonize the intestinal tract of worms might be due either to an impaired escape from detection by the immune system or to a defective resistance to its action. It is not clear, however, that RhlR-mediated regulation of the production of toxic phenazines, as shown in the case of PA14-mediated “fast killing” of *C. elegans* (Mukherjee et al., 2017), is the reason the *ΔrhlR* mutant is so dramatically impaired in the *C. elegans “*slow killing” assay used in our study in Fig. 2A. The major difference in the fast and slow *C. elegans* killing assay is the composition of the agar medium in which the *P. aeruginosa* is grown and on which the killing assays are performed (Mahajan-Miklos, Tan et al., 1999, Tan et al., 1999a, Tan, Rahme et al., 1999b). Fast killing is mediated by phenazines (Cezairliyan, Vinayavekhin et al., 2013, Mahajan-Miklos et al., 1999) whereas slow killing is multi-factorial (Feinbaum, Urbach et al., 2012), and PA14 mutants deficient in the production of phenazines do not exhibit a significant killing defect in the slow killing assay (Tan et al., 1999b). In contrast to *Drosophila*, no cellular host defense has been detected in *C. elegans* and is unlikely to be involved in the immune response to intestinal infection. The use of *ΔrhlR* mutants will open the way to the identification of the relevant host defense systems that are circumvented by wild-type PA14 bacteria. In any case, the lack of a significant phenotype of the *ΔrhlI* mutants in the *C. elegans* killing and intestinal proliferation assays is striking. This can be exploited in future studies to help elucidate the underlying *rhlI*-independent mechanisms involved in RhlR-mediated regulation of virulence.

### RhlR counteracts the cellular host defense by eluding detection by Tep4

Our phagocytosis and opsonization data are consistent with a model in which RhlR controls the expression of gene products that mask the site being recognized by Tep4 or a Tep4-associated protein, presumably on the cell wall. Alternatively, RhlR may actively inhibit the uptake of opsonized bacteria. The masking or inhibition of ingestion processes may be sensitive to heat, as wild-type heat-killed bacteria appeared to be more efficiently taken up by hemocytes than live ones (compare median values for PA14WT in Fig. 5A to those in 5B). Alternatively, the processes may be unstable and require permanent maintenance that can no longer be achieved when the bacterial cells are killed.

Insect thioester-containing proteins belong to the complement family, and have been shown to be involved in the opsonization of bacteria in mosquitoes. In *Drosophila*, Tep2 has been reported to be required for the uptake of *Escherichia coli*, a Gram-negative bacterium, by cultured S2 cells (Stroschein-Stevenson et al., 2006), a finding confirmed *in vivo* (Shokal, Kopydlowski et al., 2017). In contrast, we find no involvement of Tep2 in our *in vivo* intestinal infection model with *P. aeruginosa* but do detect a requirement for Tep4 in phagocytosis and opsonization assays. Given that the structure of mosquito thioester containing protein 1 is similar to that of complement family C3, a well-described opsonin, our data are compatible with a model of direct opsonization of bacteria by Tep4.

### A host factor plays opposite roles in host defense against the same pathogen according to the infection route

The finding that Tep4 plays a protective function in the intestinal infection model whereas it is detrimental in the case of a direct systemic infection is paradoxical. This may actually represent two faces of the same phenomenon. PA14 may have developed a stealth strategy to avoid detection by the immune system of living organisms and thus may actively hide any features that might reveal its presence. We propose here that RhlR plays a critical role in a program that renders PA14 furtive, in keeping with our finding that bacteria likely need to be alive to escape phagocytosis efficiently (Fig. 5). As a result of RhlR action, only a few sites would be available on the surface of the wild-type bacteria for Tep4 direct or indirect binding. There, Tep4 would mediate opsonization and then subsequent phagocytosis of the bacteria. Our data in the systemic infection model are compatible with the possibility that Tep4 competes for these sites with Pattern Recognition Receptors (PRRs) that activate the prophenol oxidase cascade since PO activation occurs only in the *Tep4* mutant background. It is likely that small peptidoglycan (PGN) fragments released by proliferating bacteria represent a major trigger of the IMD pathway in addition to large PGN fragments directly sensed by PGRP-LC, thereby accounting for the apparent normal expression of *Diptericin* when flies are challenged with injected PA14. In contrast, we have previously established that some fungi and Gram-positive bacteria trigger the phenol oxidase cascade through defined PRRs (Matskevich, Quintin et al., 2010). The situation is less clear as regards Gram-negative bacteria. While the original characterization of PGRP-LE suggested that it triggers the phenol oxidase activation cascade (Takehana, Katsuyama et al., 2002), and acts non cell-autonomously (Takehana, Yano et al., 2004), subsequent studies have documented a role for PGRP-LE as an intracellular sensor for PGN fragments (Bosco-Drayon, Poidevin et al., 2012, Ferrandon et al., 2007, Yano, Mita et al., 2008). Thus, further work will be required to identify whether Tep4 actually competes with PRRs in the detection of pathogens.

Our results further suggest that the cellular immune response is a relevant defense when a few bacteria enter the hemocoel after escaping from the digestive tract in the intestinal infection model, but not in the septic injury model. Conversely, melanization mediated by activated phenol oxidase appears to be somewhat effective after injection but not ingestion of PA14.

A major challenge will be to establish how RhlR influences the surface properties of PA14 or alternatively inhibits the uptake of opsonized bacteria.

Finally, *ΔrhlR* mutants exhibit reduced dissemination capacities in a rodent lung infection model when compared to *ΔrhlI* or wild-type PA14(Mukherjee et al., 2017). By analogy to our findings in the *Drosophila* intestinal infection model, it will be interesting to determine whether the complement system restricts the systemic escape of *ΔrhlR* mutants from the mouse lung into the periphery.

## MATERIALS AND METHODS

Many methods employed in this study have been described in detail in Haller *et al.* (2014).

### *C. elegans* killing and intestinal accumulation assays

The “slow-killing” of *C. elegans* by *P. aeruginosa* was monitored using automated image analysis as previously described (McEwan et al., 2016, Stroustrup, Ulmschneider et al., 2013). To monitor the accumulation of *P. aeruginosa* in the *C. elegans* intestine, wild-type (N2) animals were used for all experiments. Worms were reared on non-pathogenic *E. coli OP50* on nematode growth media at 25°C. Synchronized L4 worms (fourth larval stage) were transferred to slow kill (SK) nematode growth media agar plates containing a lawn of *P. aeruginosa* PA14::GFP strains. Post infection at 24 and 48 hours approximately 20 worms were picked onto a 2% agar pad that contain the paralyzing agent levamisole (1mM). The worms were imaged in the GFP channel using a Zeiss Apotome microscope with the same exposure time for all the worms on wild type PA14 and the *ΔrhlR* and *ΔrhlI* mutants. Post acquisition the images were processed using ImageJ software and the area and fluorescence intensity was measured. The relative fluorescence intensity is plotted and a non-parametric Mann-Whitney test was used to determine statistical significance.

### Opsonization assay of live bacteria

Overnight cultures of PA14, *ΔrhlR*, and *ΔRhlI* mutants were concentrated to OD10 in PBS. Twenty third instar larvae were bled in 150 µL of bacterial resuspended in PBS at aOD of 10 and incubated at room temperature for 30 to 45 min. Samples were centrifuged at 500 rcf for 15 min and the pellet (containing larval debris) was removed. A second centrifugation was performed at 3500 rcf for 15 min to retrieve bacteria in the pellet, that was re-suspended in 10 µL PBS. A5001 or *tep4* third instar larvae were injected with 32.2 nL of the live bacteria solutions, using a Nanoject apparatus (Drummont). After 60-90 min of incubation, one larva was bled in each well of an 8-well pattern microscopy slide that contained PBS. The cells were left to settle to the bottom for 30 min and were then fixed in 4 % paraformaldehyde for 15 min, in a humid chamber. The samples were washed twice in PBS and they were stained with a 1/500 diluted rabbit antiserum against PA14 in a PBS solution with 2 % BSA for 2 hours at room temperature. The cells were incubated with a FITC-labeled goat anti-rabbit secondary antibody (Invitrogen) in a PBS solution with 2 % BSA for 2 hours at room temperature. After a 20 min permeabilization step in a PBS solution with 0.1 % Triton X-100 and 2 % BSA, a second round of staining with a 1/500 diluted rabbit antiserum against PA14 in a PBS solution with 0.1 % Triton X-100 and 2 % BSA was performed for 2 hours at room temperature. The samples were then incubated with a Cy3-labeled goat anti-rabbit secondary antibody (Invitrogen) in a PBS solution with 0.1 % Triton X-100 and 2 % BSA for 2 hours at room temperature. The slides were mounted in Vectashield with DAPI (Vector Laboratories) and analyzed using a Zeiss Axioskope 2 fluorescent microscope. 40 to 50 cells per larva were analyzed: the number of red fluorescent bacteria that were not green fluorescent was counted for each DAPI-positive hemocyte, and the phagocytosis index was calculated (% of phagocytes containing at least 1 only-green bacterium)×(mean number of only-green bacteria per positive cell). We used the nonparametric Mann-Whitney test for statistical analysis.

### Phenol oxidase cleavage assay

The procedure was performed as described (Leclerc, Pelte et al., 2006), except that hemolymph loads were not adjusted by measuring the protein content of the extracted hemolymph. The antibody used has been generated by Dr. H. M. Müller against *Anopheles* phenol oxidases (Muller, Dimopoulos et al., 1999). The ratio of cleaved to noncleaved form was performed by densitometry scanning.

## Statistical analysis

All statistical analysis were performed on Graphpad Prism version 5 (Graphpad software Inc., San Diego, CA). Details are indicated in the legend of each figure.

## ACKNOWLEDGEMENTS

We are grateful to J. Nguyen, to Gaétan Caravello, and D. McEwan for help with some experiments, and to W. M. Yamba and J. Bourdeau for expert technical help. We thank Dr. A. Filloux and Dr. B. Bassler for the gift of PA14 mutant strains, the Bloomington Stock Center for *Drosophila* stocks. Dr. S. Niehus provided valuable advice. We thank Dr. B. Bassler for sharing her work with us prior to publication. This work has been funded by CNRS, University of Strasbourg, Fondation pour la Recherche Médicale (Equipe FRM to D.F.), Agence Nationale de la Recherche (DROSOGUT, ANR-11-EQPX-0022) and US NIH grant R01 AI085581 awarded to F.M.A.

## AUTHOR CONTRIBUTIONS

SH, AF, SL, and DF conceived the *Drosophila* experiments and analyzed the data. SH performed the majority of these experiments, with later work performed by AF with an input from SL; SS performed the precursor experiments that led to this work. Work on the septic injury model was performed by JC, with some help from ZL. *ΔlasR*, ΔlasI, ΔrhlR and *ΔrhlI* mutants were constructed by ED and SY, except when indicated otherwise. AH and FMA conceived the *C. elegans* experiments and analyzed the data, which were obtained by AH. SH, DF, and FMA wrote the manuscript, with inputs from other co-authors.

## REFERENCES

Anderson RM, Zimprich CA, Rust L(1999) A second operator is involved in Pseudomonas aeruginosa elastase (lasB) activation. J Bacteriol 181: 6264–70

Arefin B, Kucerova L, Dobes P, Markus R, Strnad H, Wang Z, Hyrsl P, Zurovec M, Theopold U. (2014) Genome-wide transcriptional analysis of Drosophila larvae infected by entomopathogenic nematodes shows involvement of complement, recognition and extracellular matrix proteins. J Innate Immun 6: 192–204

Avet-Rochex A, Bergeret E, Attree I, Meister M, Fauvarque MO (2005) Suppression of Drosophila cellular immunity by directed expression of the ExoS toxin GAP domain of Pseudomonas aeruginosa. Cell Microbiol 7: 799–810

Ayyaz A, Li H, Jasper H (2015) Haemocytes control stem cell activity in the Drosophila intestine. Nat Cell Biol 17: 736–48

Batz T, Forster D, Luschnig S (2014) The transmembrane protein Macroglobulin complement-related is essential for septate junction formation and epithelial barrier function in Drosophila. Development 141: 899–908

Bier E, Guichard A (2012) Deconstructing host-pathogen interactions in Drosophila. Dis Model Mech 5: 48–61

Binggeli O, Neyen C, Poidevin M, Lemaitre B (2014) Prophenoloxidase activation is required for survival to microbial infections in Drosophila. PLoS Pathog 10: e1004067

Bosco-Drayon V, Poidevin M, Boneca IG, Narbonne-Reveau K, Royet J, Charroux B (2012) Peptidoglycan sensing by the receptor PGRP-LE in the Drosophila gut induces immune responses to infectious bacteria and tolerance to microbiota. Cell Host Microbe 12: 153–65

Bou Aoun R, Hetru C, Troxler L, Doucet D, Ferrandon D, Matt N (2011) Analysis of thioester-containing proteins during the innate immune response of Drosophila melanogaster. J Innate Immun 3: 52–64

Buchon N, Silverman N, Cherry S (2014) Immunity in Drosophila melanogaster—from microbial recognition to whole-organism physiology. Nat Rev Immunol 14: 796–810

Campbell J, Lin Q, Geske GD, Blackwell HE (2009) New and unexpected insights into the modulation of LuxR-type quorum sensing by cyclic dipeptides. ACS chemical biology 4: 1051–9

Cezairliyan B, Vinayavekhin N, Grenfell-Lee D, Yuen GJ, Saghatelian A, Ausubel FM (2013) Identification of Pseudomonas aeruginosa phenazines that kill Caenorhabditis elegans. PLoS Pathog 9: e1003101

Coggan KA, Wolfgang MC (2012) Global regulatory pathways and cross-talk control pseudomonas aeruginosa environmental lifestyle and virulence phenotype. Current issues in molecular biology 14: 47–70

Dudzic JP, Kondo S, Ueda R, Bergman CM, Lemaitre B (2015) Drosophila innate immunity: regional and functional specialization of prophenoloxidases. BMC Biol 13: 81

El Chamy L, Matt N, Reichhart JM (2017) Advances in Myeloid-Like Cell Origins and Functions in the Model Organism Drosophila melanogaster. Microbiol Spectr 5

Elrod-Erickson M, Mishra S, Schneider D (2000) Interactions between the cellular and humoral immune responses in *Drosophila*. Curr Biol 10: 781–4

Feinbaum RL, Urbach JM, Liberati NT, Djonovic S, Adonizio A, Carvunis AR, Ausubel FM (2012) Genome-wide identification of Pseudomonas aeruginosa virulence-related genes using a Caenorhabditis elegans infection model. PLoS Pathog 8: e1002813

Ferrandon D (2013) The complementary facets of epithelial host defenses in the genetic model organism Drosophila melanogaster: from resistance to resilience. Curr Opin Immunol 25: 59–70

Ferrandon D, Imler JL, Hetru C, Hoffmann JA (2007) The Drosophila systemic immune response: sensing and signalling during bacterial and fungal infections. Nat Rev Immunol 7: 862–74

Gambello MJ, Kaye S, Iglewski BH (1993) LasR of Pseudomonas aeruginosa is a transcriptional activator of the alkaline protease gene (apr) and an enhancer of exotoxin A expression. Infect Immun 61: 1180–4

Ganesan S, Aggarwal K, Paquette N, Silverman N (2010) NF-kappaB/Rel Proteins and the Humoral Immune Responses of Drosophila melanogaster. Curr Top Microbiol Immunol

Hall S, Bone C, Oshima K, Zhang L, McGraw M, Lucas B, Fehon RG, Ward REt (2014) Macroglobulin complement-related encodes a protein required for septate junction organization and paracellular barrier function in Drosophila. Development 141: 889–98

Haller S, Limmer S, Ferrandon D (2014) Assessing Pseudomonas virulence with a nonmammalian host: Drosophila melanogaster. Methods Mol Biol 1149: 723–40

Holden MT, Ram Chh.bra S, de Nys R, Stead P, Bainton NJ, Hill PJ, Manefield M, Kumar N, Labatte M, England D, Rice S, Givskov M, Salmond GP, Stewart GS, Bycroft BW, Kjelleberg S, Williams P (1999) Quorum-sensing cross talk: isolation and chemical characterization of cyclic dipeptides from Pseudomonas aeruginosa and other gram-negative bacteria. Mol Microbiol 33: 1254–66

Hoyland-Kroghsbo NM, Paczkowski J, Mukherjee S, Broniewski J, Westra E, Bondy-Denomy J, Bassler BL (2017) Quorum sensing controls the Pseudomonas aeruginosa CRISPR-Cas adaptive immune system. Proc Natl Acad Sci U S A 114: 131–135

Igboin CO, Griffen AL, Leys EJ (2012) The Drosophila melanogaster host model. Journal of oral microbiology 4

Irazoqui JE, Urbach JM, Ausubel FM (2010) Evolution of host innate defence: insights from Caenorhabditis elegans and primitive invertebrates. Nat Rev Immunol 10: 47–58

Jimenez PN, Koch G, Thompson JA, Xavier KB, Cool RH, Quax WJ (2012) The multiple signaling systems regulating virulence in Pseudomonas aeruginosa. Microbiology and molecular biology reviews : MMBR 76: 46–65

Kocks C, Cho JH, Nehme N, Ulvila J, Pearson AM, Meister M, Strom C, Conto SL, Hetru C, Stuart LM, Stehle T, Hoffmann JA, Reichhart JM, Ferrandon D, Ramet M, Ezekowitz RA (2005) Eater, a transmembrane protein mediating phagocytosis of bacterial pathogens in Drosophila. Cell 123: 335–46

Latifi A, Winson MK, Foglino M, Bycroft BW, Stewart GS, Lazdunski A, Williams P (1995) Multiple homologues of LuxR and LuxI control expression of virulence determinants and secondary metabolites through quorum sensing in Pseudomonas aeruginosa PAO1. Mol Microbiol 17: 333–43

Leclerc V, Pelte N, El Cha. y L, Martinelli C, Ligoxygakis P, Hoffmann JA, Reichhart JM (2006) Prophenoloxidase activation is not required for survival to microbial infections in Drosophila. EMBO Rep 7: 231–5

Ledgham F, Ventre I, Soscia C, Foglino M, Sturgis JN, Lazdunski A (2003) Interactions of the quorum sensing regulator QscR: interaction with itself and the other regulators of Pseudomonas aeruginosa LasR and RhlR. Mol Microbiol 48: 199–210

Lemaitre B, Hoffmann J (2007) The Host Defense of Drosophila melanogaster. Annu Rev Immunol 25: 697–743

Levashina EA, Moita LF, Blandin S, Vriend G, Lagueux M, Kafatos FC (2001) Conserved role of a complement-like protein in phagocytosis revealed by dsRNA knockout in cultured cells of the mosquito, Anopheles gambiae. Cell 104: 709–18

Limmer S, Haller S, Drenkard E, Lee J, Yu S, Kocks C, Ausubel FM, Ferrandon D (2011a) Pseudomonas aeruginosa RhlR is required to neutralize the cellular immune response in a Drosophila melanogaster oral infection model. Proc Natl Acad Sci U S A 108: 17378–83

Limmer S, Quintin J, Hetru C, Ferrandon D (2011b) Virulence on the fly: Drosophila melanogaster as a model genetic organism to decipher host-pathogen interactions. Current Drug Targets 12: 978–999

Lin L, Rodrigues F, Kary C, Contet A, Logan M, Baxter RHG, Wood W, Baehrecke EH (2017) Complement-Related Regulates Autophagy in Neighboring Cells. Cell 170: 158–171 e8

Mahajan-Miklos S, Tan MW, Rahme LG, Ausubel FM (1999) Molecular mechanisms of bacterial virulence elucidated using a Pseudomonas aeruginosa-Caenorhabditis elegans pathogenesis model. Cell 96: 47–56

Matskevich AA, Quintin J, Ferrandon D (2010) The Drosophila PRR GNBP3 assembles effector complexes involved in antifungal defenses independently of its Toll-pathway activation function. Eur J Immunol 40: 1244–54

McEwan DL, Feinbaum RL, Stroustrup N, Haas W, Conery AL, Anselmo A, Sadreyev R, Ausubel FM (2016) Tribbles ortholog NIPI-3 and bZIP transcription factor CEBP-1 regulate a Caenorhabditis elegans intestinal immune surveillance pathway. BMC Biol 14: 105

Medina G, Juarez K, Valderrama B, Soberon-Chavez G (2003) Mechanism of Pseudomonas aeruginosa RhlR transcriptional regulation of the rhlAB promoter. J Bacteriol 185: 5976–83

Mukherjee S, Moustafa D, Smith CD, Goldberg JB, Bassler BL (2017) The RhlR quorum-sensing receptor controls Pseudomonas aeruginosa pathogenesis and biofilm development independently of its canonical homoserine lactone autoinducer. PLoS Pathog 13: e1006504

Muller HM, Dimopoulos G, Blass C, Kafatos FC (1999) A hemocyte-like cell line established from the malaria vector Anopheles gambiae expresses six prophenoloxidase genes. J Biol Chem 274: 11727–35

Nehme NT, Liegeois S, Kele B, Giammarinaro P, Pradel E, Hoffmann JA, Ewbank JJ, Ferrandon D (2007) A Model of Bacterial Intestinal Infections in Drosophila melanogaster. PLoS Pathog 3: e173

Pean CB, Dionne MS (2014) Intracellular infections in Drosophila melanogaster: host defense and mechanisms of pathogenesis. Dev Comp Immunol 42: 57–66

Pesci EC, Pearson JP, Seed PC, Iglewski BH (1997) Regulation of las and rhl quorum sensing in Pseudomonas aeruginosa. J Bacteriol 179: 3127–32

Pukkila-Worley R, Ausubel FM (2012) Immune defense mechanisms in the Caenorhabditis elegans intestinal epithelium. Curr Opin Immunol 24: 3–9

Schuster M, Lostroh CP, Ogi T, Greenberg EP (2003) Identification, timing, and signal specificity of Pseudomonas aeruginosa quorum-controlled genes: a transcriptome analysis. J Bacteriol 185: 2066–79

Schuster M, Sexton DJ, Diggle SP, Greenberg EP (2013) Acyl-homoserine lactone quorum sensing: from evolution to application. Annu Rev Microbiol 67: 43–63

Seed PC, Passador L, Iglewski BH (1995) Activation of the Pseudomonas aeruginosa lasI gene by LasR and the Pseudomonas autoinducer PAI: an autoinduction regulatory hierarchy. J Bacteriol 177: 654–9

Shokal U, Eleftherianos I (2017) Thioester-Containing Protein-4 Regulates the Drosophila Immune Signaling and Function against the Pathogen Photorhabdus. J Innate Immun 9: 83–93

Shokal U, Kopydlowski H, Eleftherianos I (2017) The distinct function of Tep2 and Tep6 in the immune defense of Drosophila melanogaster against the pathogen Photorhabdus. Virulence: 1–15

Stroschein-Stevenson SL, Foley E, O’Farrell PH, Johnson AD (2006) Identification of Drosophila gene products required for phagocytosis of Candida albicans. PLoS Biol 4: e4

Stroustrup N, Ulmschneider BE, Nash ZM, Lopez-Moyado IF, Apfeld J, Fontana W (2013) The Caenorhabditis elegans Lifespan Machine. Nat Methods 10: 665–70

Takehana A, Katsuyama T, Yano T, Oshima Y, Takada H, Aigaki T, Kurata S (2002) Overexpression of a pattern-recognition receptor, peptidoglycan-recognition protein-LE, activates imd/relish-mediated antibacterial defense and the prophenoloxidase cascade in Drosophila larvae. Proc Natl Acad Sci U S A 99: 13705–10

Takehana A, Yano T, Mita S, Kotani A, Oshima Y, Kurata S (2004) Peptidoglycan recognition protein (PGRP)-LE and PGRP-LC act synergistically in Drosophila immunity. Embo J 23: 4690–700

Tan MW, Mahajan-Miklos S, Ausubel FM (1999a) Killing of Caenorhabditis elegans by Pseudomonas aeruginosa used to model mammalian bacterial pathogenesis. Proc Natl Acad Sci U S A 96: 715–20

Tan MW, Rahme LG, Sternberg JA, Tompkins RG, Ausubel FM (1999b) Pseudomonas aeruginosa killing of Caenorhabditis elegans used to identify P. aeruginosa virulence factors. Proc Natl Acad Sci U S A 96: 2408–13

Wagner VE, Bushnell D, Passador L, Brooks AI, Iglewski BH (2003) Microarray analysis of Pseudomonas aeruginosa quorum-sensing regulons: effects of growth phase and environment. J Bacteriol 185: 2080–95

Williams P, Camara M (2009) Quorum sensing and environmental adaptation in Pseudomonas aeruginosa: a tale of regulatory networks and multifunctional signal molecules. Curr Opin Microbiol 12: 182–91

Yano T, Mita S, Ohmori H, Oshima Y, Fujimoto Y, Ueda R, Takada H, Goldman WE, Fukase K, Silverman N, Yoshimori T, Kurata S (2008) Autophagic control of listeria through intracellular innate immune recognition in drosophila. Nat Immunol 9: 908–16

Zugasti O, Bose N, Squiban B, Belougne J, Kurz CL, Schroeder FC, Pujol N, Ewbank JJ (2014) Activation of a G protein-coupled receptor by its endogenous ligand triggers the innate immune response of Caenorhabditis elegans. Nat Immunol 15: 833–8

